## Supplementary material for "Adaptation of *Bacillus thuringiensis* to plant colonization affects differentiation and toxicity": Table S1 and Fig S1 to S7

**Table S1** Information of strains, plasmids and oligos used in this study

| | Description | Reference |
|---|---|---|
| **Strains** | | |
| *Bacillus thuringiensis* 407 cry- (Bt407) | The acrystalliferous *B. thuringiensis* 407 strain obtained by high temperature process | (1) |
| Bt407 GFP | Bt407 transformed with pTB603 | This study |
| Bt407 mKate | Bt407 transformed with pTB604 | This study |
| Bt407 EvoA GFP | Bt407 evolved isolate from lineage A transformed with pTB603 | This study |
| Bt407 EvoC GFP | Bt407 evolved isolate from lineage C transformed with pTB603 | This study |
| Bt407 EvoDGFP | Bt407 evolved isolate from lineage D transformed with pTB603 | This study |
| Bt407 EvoE GFP | Bt407 evolved isolate from lineage E transformed with pTB603 | This study |
| Bt407 EvoF GFP | Bt407 evolved isolate from lineage F transformed with pTB603 | This study |
| Bt407 $rho^{Glu54stop}$ | Bt407 introduced a mutation of $rho^{Glu54stop}$ | This study |
| *Escherichia coli* XL-1 Blue | *E. coli* K-12 derivatives used for cloning procedures | (2) |
| **Plasmids** | | |
| pMAD | Used for homologous recombination in gram positive strains ($Amp^R$ and $Ery^R$) | (3) |
| pMAD (I-SceI) | pMAD containing I-SceI restriction digestion site ($Amp^R$ and $Ery^R$) | (4) |
| pBKJ223 | Harboring I-SceI enzyme for restriction digestion ($Amp^R$ and $Tet^R$) | (5) |
| **Oligos** | | |
| oYL29 | Forward: amplify the fragment from Rho mutation CCG GAA TTC TTC AAT ATT ACG TGC CGC T | This study |
| oYL30 | Reverse: amplify the fragment from E contain rho mutation CGC GGA TCC TGG TGA ACT ATT ACG TGG T | This study |
| oYL33 | Forward: amplify the fragment from SigE mutation CCG GAA TTC CGC TTA TCT CTC CAC TTT | This study |
| oYL34 | Reverse: amplify the fragment from SigE mutation CGC GGA TCC ACC CAT TAA CAA AAA CAC CC | This study |
| oYL41 | Forward: amplify the fragment from pMAD CGTCATATGGATCCGATATC | This study |
| oYL42 | Reverse: amplify the fragment from pMAD ATGGCATGCATCGATAGATC | This study |
| oYL45 | Forward: check the successful integration of pMAD with rho mutation into chromosome of Bt GTTCCATATTTCCAGTTCC | This study |
| oYL46 | Reverse: check the successful integration of pMAD with rho mutation into chromosome of Bt CCACAAAACGAAGCTGAA | This study |
| oYL47 | Forward: check the successful integration of pMAD with SigE mutation into chromosome of Bt AATAATCATCGGCACAGCA | This study |
| oYL48 | Reverse: check the successful integration of pMAD with SigE mutation into chromosome of Bt CACGAAAATACACTATGACC | This study |
| oYL49 | Forward: for sequencing the insert of MCS sites of pMAD TCTATCGATGCATGCCAT | This study |
| oYL50 | Reverse: for sequencing the insert of MCS sites of pMAD AGAATCATAATGGGGAAGG | This study |

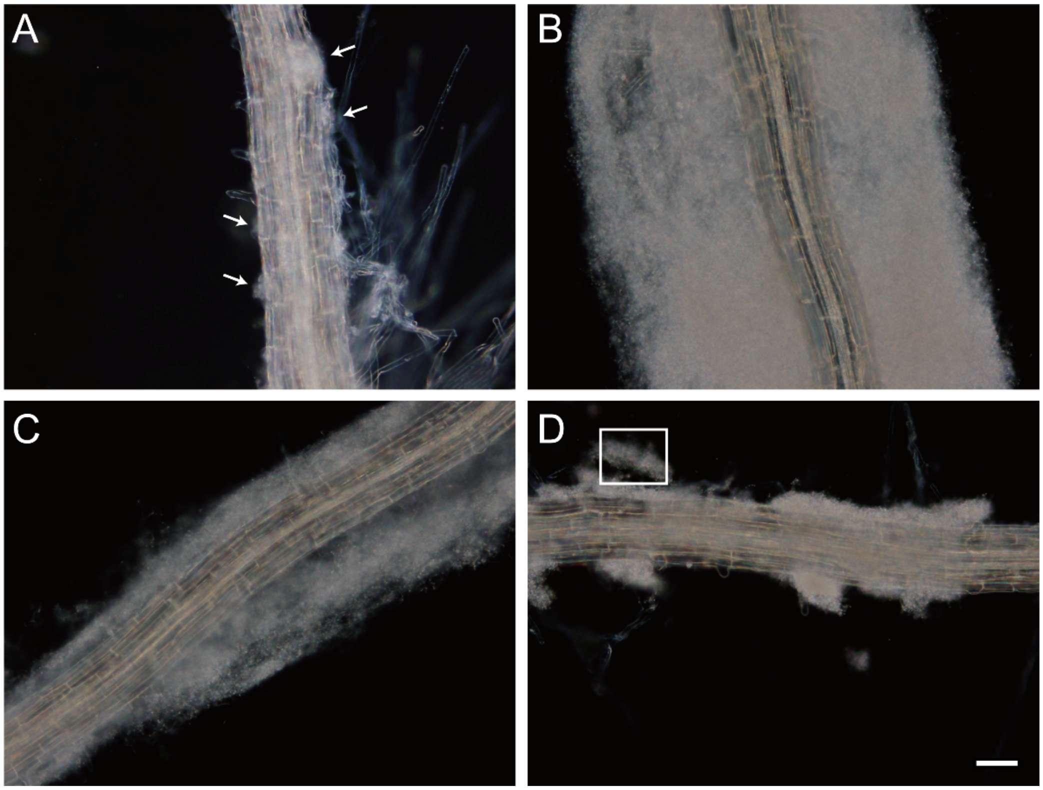

**Fig. S1 Evolved root associated biofilms of Bt407. (A)** Bt407 ancestor formed microcolonies on root of *A. thaliana*. White arrows indicate typical microcolonies. **(B)** Evolved isolate E formed extensive biofilms (even up to several times thicker than the root). **c,** Most of root associated biofilms were lost due to the shear force during the transfers, leaving microcolonies colonizing the seedlings. Images showed before **(C)** and after **(D)** wash procedure. Bacterial aggregates (indicated by white box) were dispersed from root by shear force. All images are representative of three independent cultures. Scale bar indicates 200 μm.

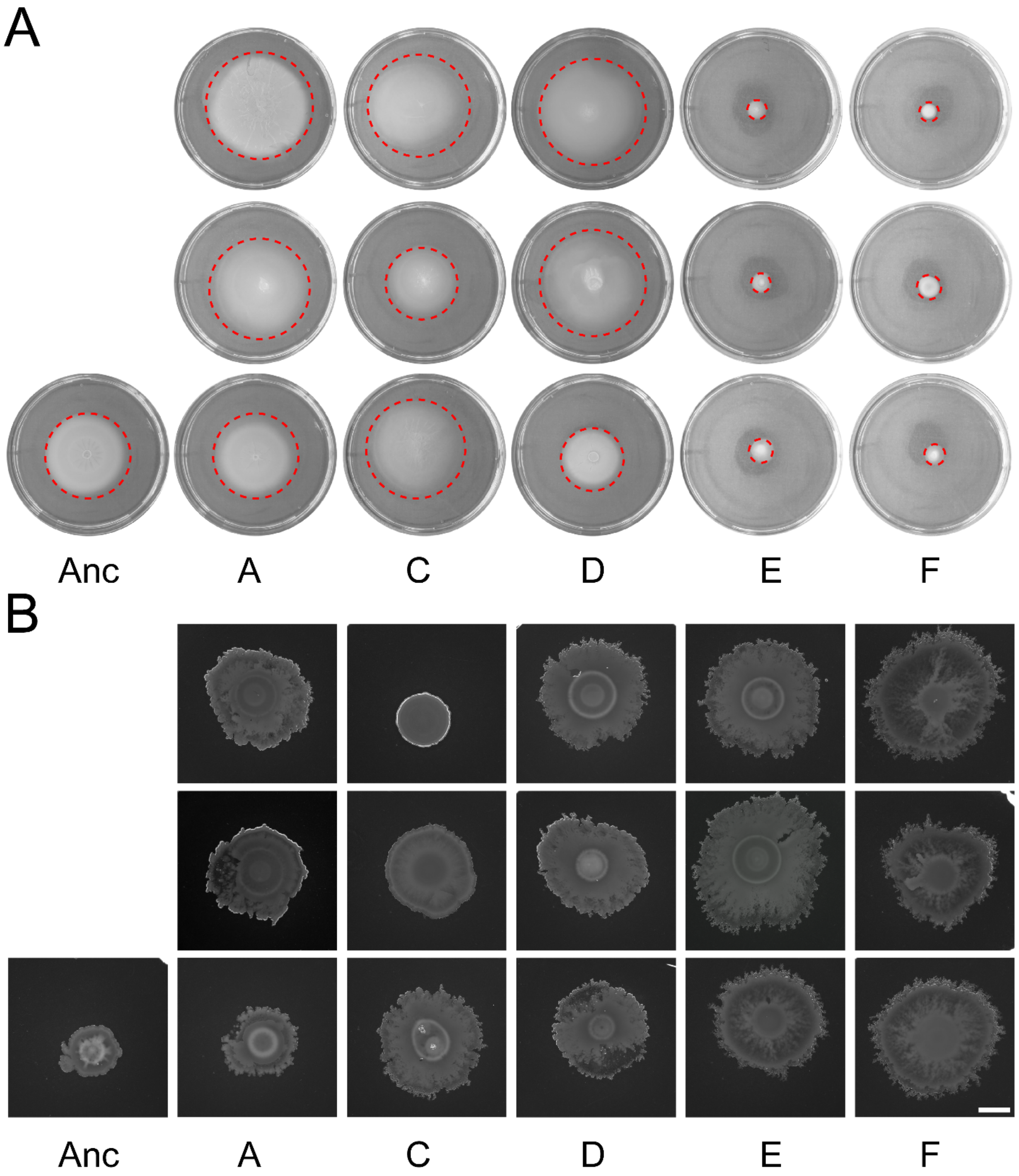


**Fig. S2 Swimming and swarming motility profiles of all isolates from each lineage. (A)** Representative images of swimming radius of three isolates from all evolved lineages. **(B)** Representative images of swarming colonies. Scale bar indicates 1 cm. Images are representatives from at least three biologically independent replicates.

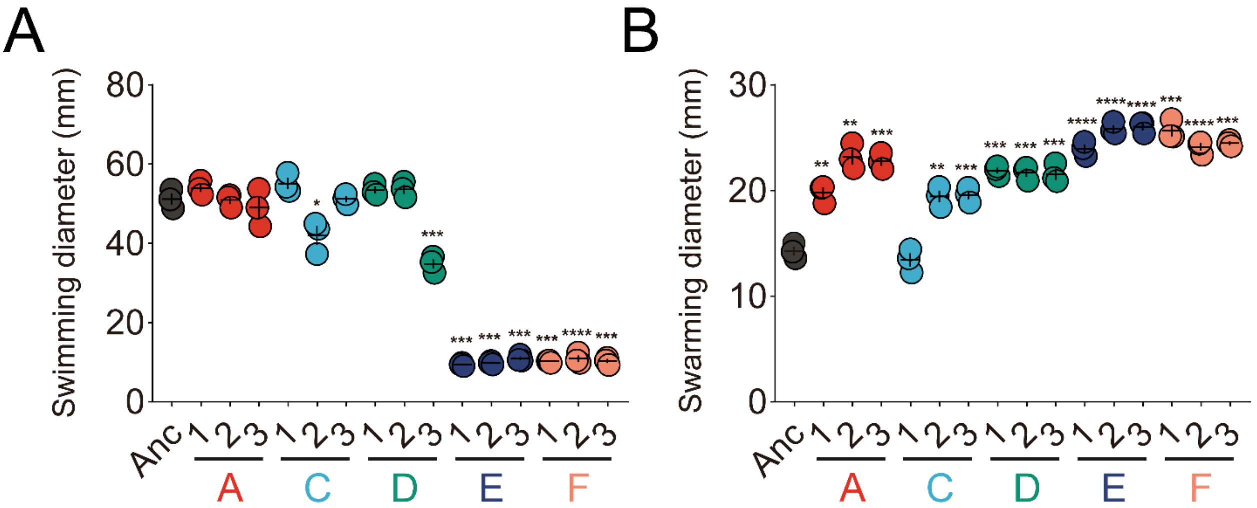


**Fig. S3 Quantification of swimming and swarming motility of all isolates from each lineage. (A)** Swimming motility quantification on plates. **(B)** Swarming motility quantification on plates. Error bars indicate standard error of means. Statistical analysis was conducted for motility diameter of evolved isolates versus ancestor using Student's *t* test with Welch correction. *p* values are demonstrated as: *p* * < 0.05, ** *p* < 0.01, *** *p* < 0.001, **** *p* < 0.0001.

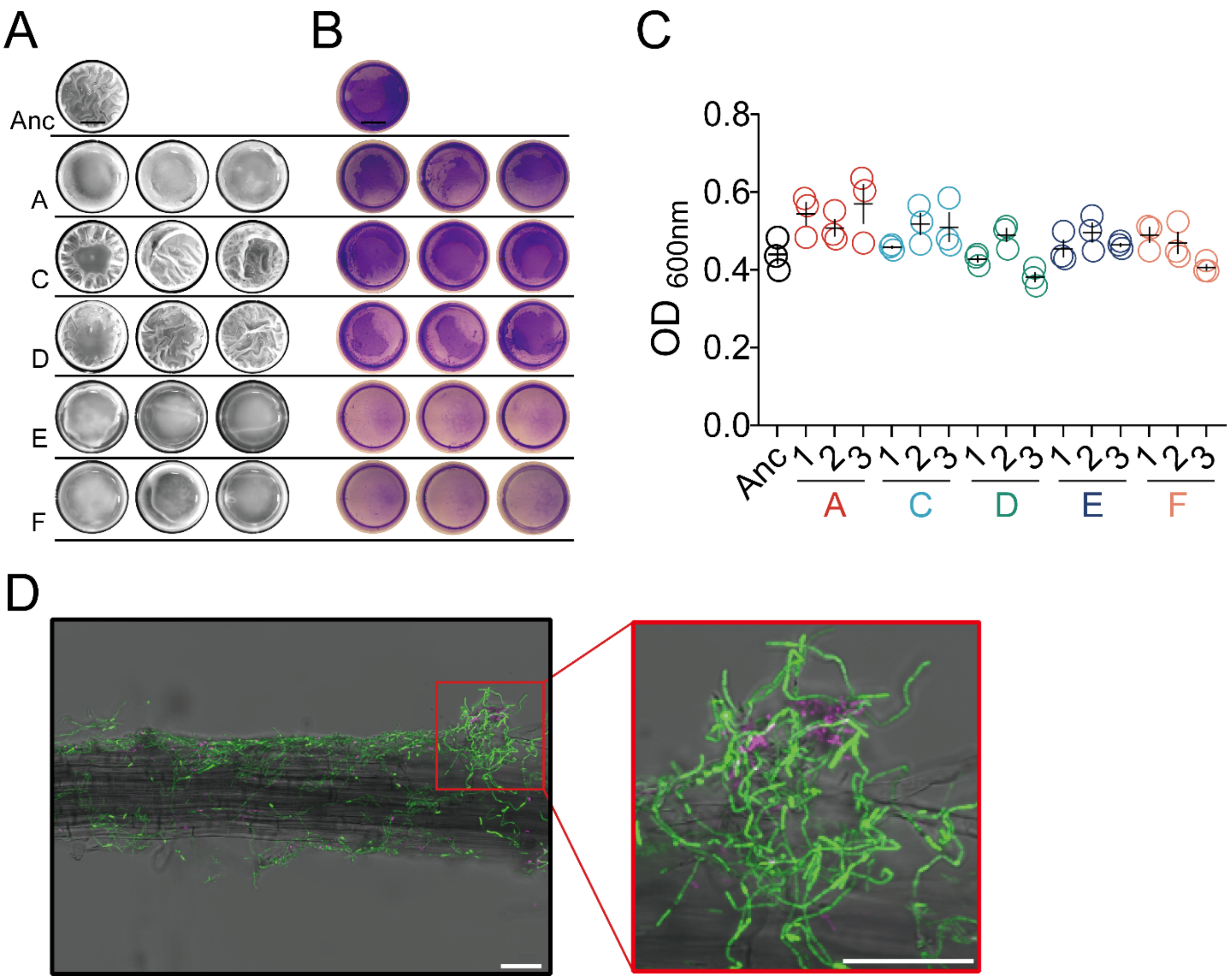


**Fig. S4 Biofilm formation properties of all strains and evolved strain E exhibited distinct cell morphology when competitively forming biofilms on the root surface of *A. thaliana*. (A)** Pellicle formation and **(B)** submerged attached biofilms were examined in 24-well microtiter plates. Images were representatives from three biologically independent replicates. Scale bar indicates 3 mm. **(C)** Total growth was measured as $OD_{600nm}$ of each sample during submerged biofilm measurement. The central values (horizontal lines) represent the mean (n=3 biologically independent samples) and the error bars represent SEM (standard error of the mean). No significant difference was found across all groups using unpaired two-tailed t-test with Welch's corrections. **(D)** Bt cells that carried either mKate (magenta, ancestor) or GFP (green, evolved isolate) reporter were co-cultured with *A. thaliana* seedlings at a one-to-one ratio. After 48h of incubation, root biofilms were visualized by CLSM. Bt407 ancestor and evolved isolate demonstrate different cell morphologies in root biofilms. Scale bar indicates 50 µm.

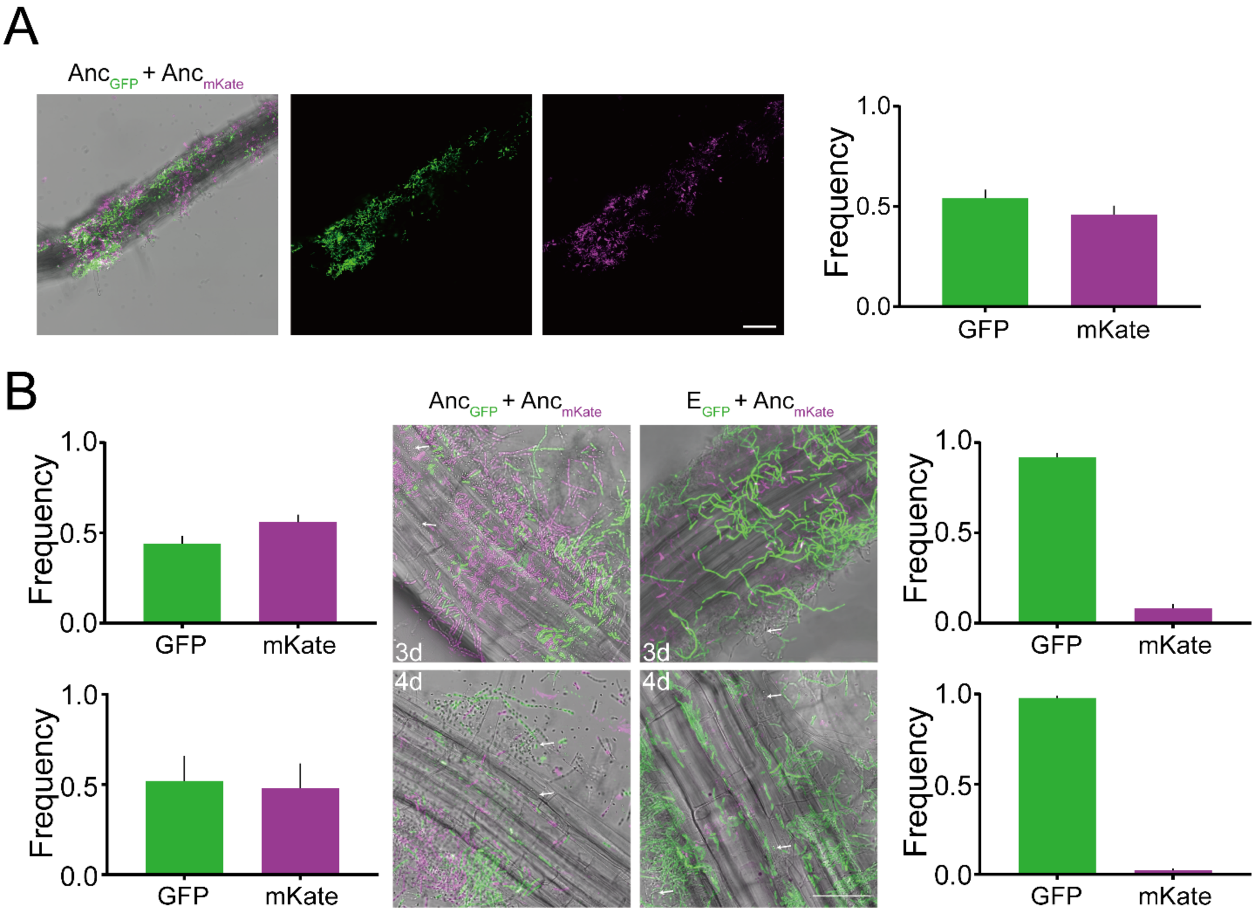


**Fig. S5 Colonization of Bt407 ancestor strains labeled with mKate and GFP at a one-to-one ratio. (A)** CLSM imaging revealed root colonization by Bt407 ancestor mixture. Images are representatives from three biologically independent plantlets images. Imaging analysis presented here revealed no significant difference between mKate and GFP signal intensity. Statistical analysis was performed by unpaired two-tailed Student's t test. Scale bars indicate 50 µm. **(B)** Bt407 ancestor and evolved strain E shows different sporulation properties associated with *A. thaliana* seedlings. Bt cells that carried mKate (magenta, ancestor) or GFP (green, evolved isolate) reporters were co-cultured with *A. thaliana* seedlings at a one-to-one ratio. After 3 and 4 days of incubation, root biofilms were visualized by CLSM. Images are representatives of three independent experiments. White arrows indicate matured spores captured by CLSM. Scale bars indicate 45 µm. Bar plots indicate fluorescent ratio between GFP and mKate ($n$=3 representing independent plantlets).

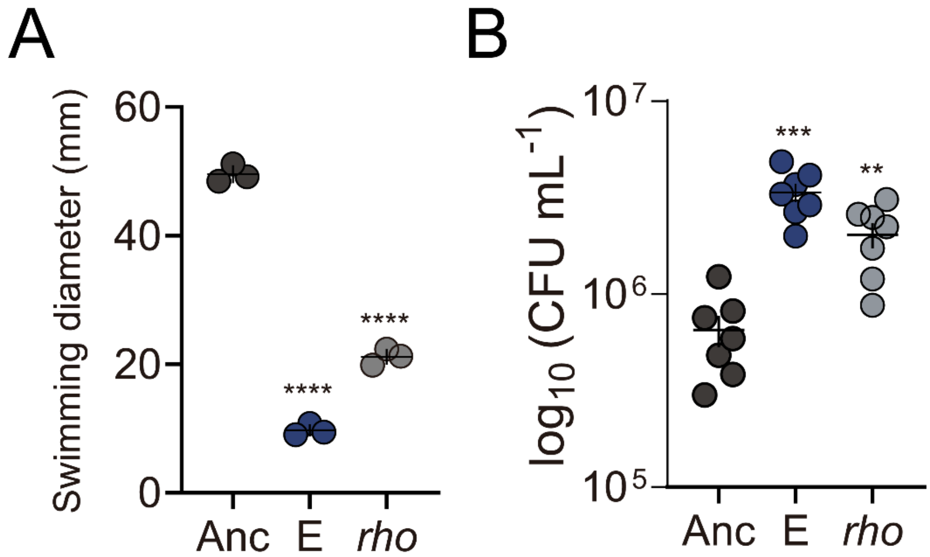


**Fig. S6 Quantification of swimming diameter of *rho*$^{Glu54stop}$ strain and aggregation size corresponding to the total number of cells observed in the largest aggregates. (A)** Swimming motility quantification on plates. **(B)** Total CFU analysis of the largest aggregates for ancestor, evolved lineage E and *rho*$^{Glu54stop}$ strain. Error bars indicate standard error of means. Statistical analysis was conducted for evolved and reconstructed strain versus ancestor using Student's *t* test with Welch correction. *p* values are demonstrated as: ** $p < 0.01$, *** $p < 0.001$, **** $p < 0.0001$.

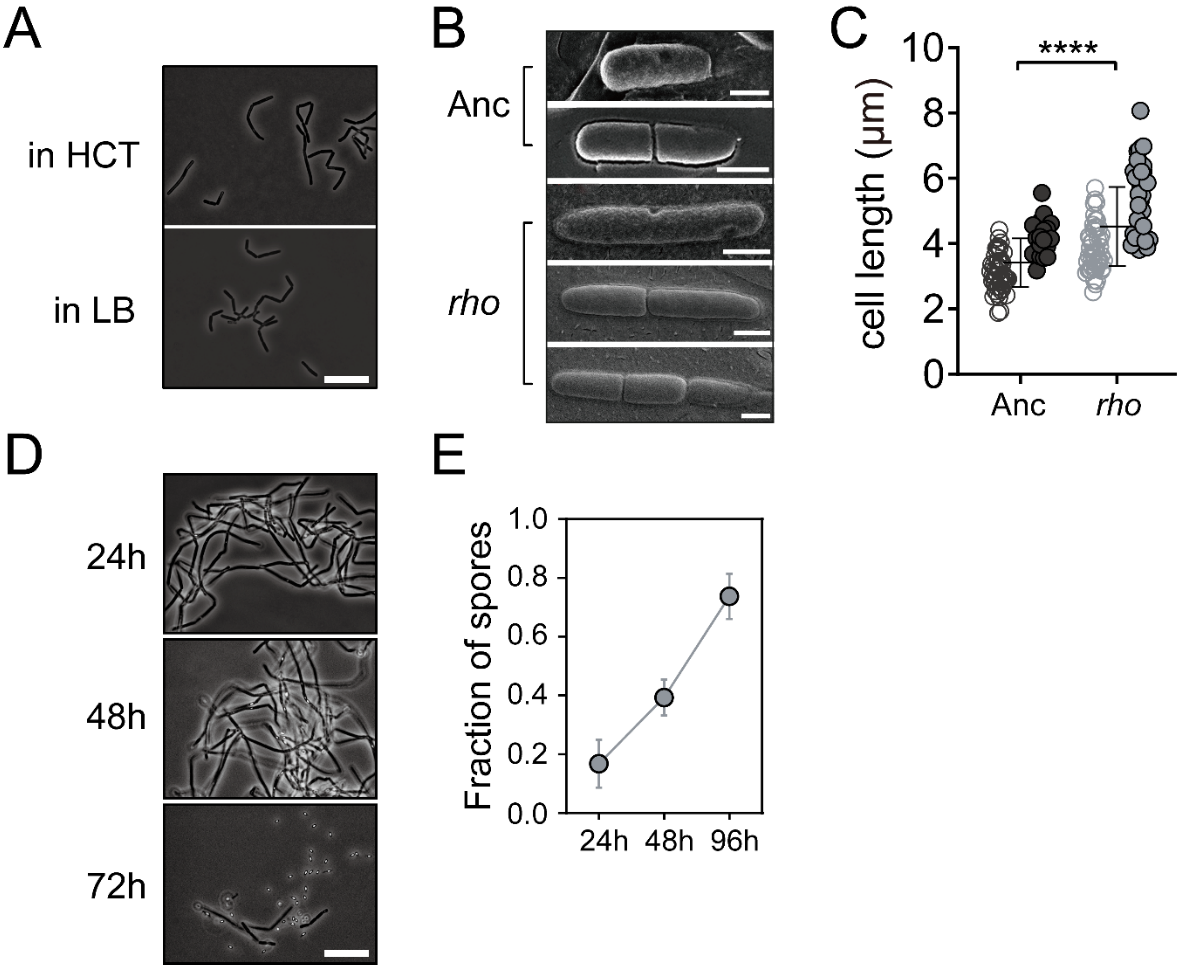


**Fig. S7 Transcriptional termination factor Rho controls cell division fate. (A)** Cell morphology was imaged under contrast phase microscopy and images representative from three biological replicates were shown. Scale bar indicates 10 μm. **(B)** Different panels are representative images of single cell and filaments consist of two or three cells captured by Cryo-SEM imaging. Scale bar indicates 1 μm. **(C)** Cell (open symbols) and chain (closed symbols) length of ancestor and $rho^{\mathrm{Glu54stop}}$ strain. Two-tailed unpaired Student's t tests were performed; **** denote $p < 0.0001$. **(D)** Microscopic observations of $rho^{\mathrm{Glu54stop}}$ strain in MSNc medium. Three independent cultures were grown and harvested at different time points. Ten microliters of vigorously vortexed cultures were observed under contrast phase microscope and representative images are shown. Scale bar indicates 10 μm. **(E)** Sporulation efficacy at three time points. Symbols represent mean of three assays and error bars represent the s.d. (standard deviation) on the mean values.
